# Substrate complexity buffers negative interactions in a synthetic microbial community of leaf litter degraders

**DOI:** 10.1101/2023.12.23.573222

**Authors:** Parmis Abdoli, Clément Vulin, Miriam Lepiz, Alexander B. Chase, Claudia Weihe, Alejandra Rodríguez-Verdugo

**Author notes:** Corresponding author: Alejandra Rodríguez-Verdugo.

## Abstract

Microbes associated with leaf litter, the top layer of soil, collectively decompose organic matter such as plant polysaccharides, and respire carbon dioxide, regulating the land-atmosphere fluxes of carbon. Therefore, it is crucial to understand the processes limiting biopolymer degradation and their influences on soil community properties. For example, it is still unclear how substrate complexity – defined as the structure of the saccharide and the amount of external processing by extracellular enzymes – influences species interactions and species coexistence. Here, we tested the hypothesis that growth on monosaccharides (i.e., xylose) promotes negative interactions through resource competition, and growth on polysaccharides (i.e., xylan) promotes neutral or positive interactions through resource partitioning or synergism among extracellular enzymes. We assembled a three-species community of leaf litter-degrading bacteria isolated from a grassland site in Southern California. In the polysaccharide xylan, pairs of species stably coexisted and grew equally in co-culture and in monoculture. Conversely, in the monosaccharide xylose, competitive exclusion and negative interactions prevailed. These pairwise dynamics remained consistent in a three-species community: all three species coexisted in xylan, while only two species coexisted in xylose. A mathematical model parameterized from single-species growth behaviors showed that in xylose these dynamics could be explained by resource competition. Instead, the resource competition model could not predict the coexistence patterns in xylan. Overall, our study shows that substrate complexity influences species interactions and patterns of coexistence in a synthetic microbial community of leaf litter degraders that can serve as a model for studying carbon cycling and climate change.

**Graphical Abstract:** 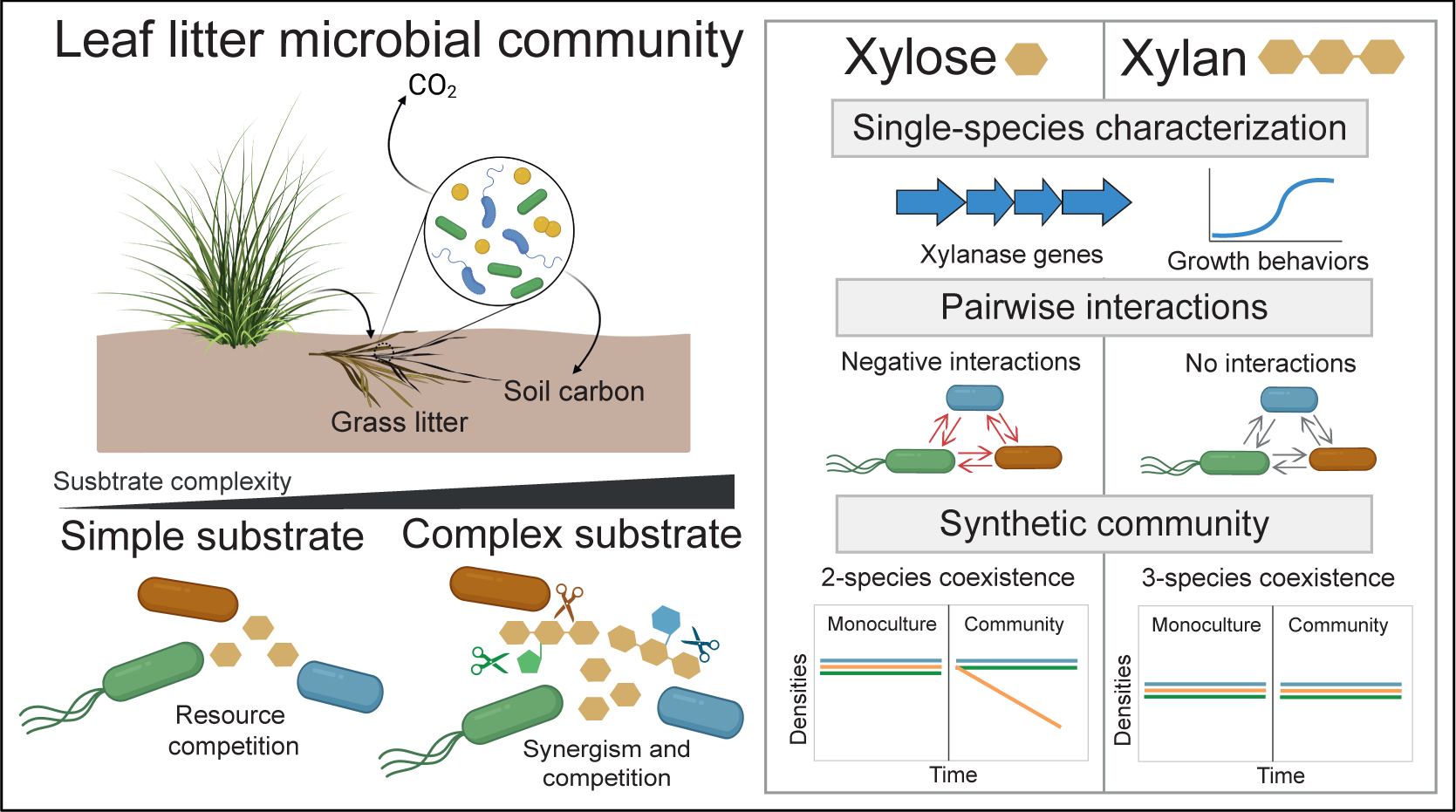

## Introduction

Microbial communities regulate biogeochemical processes, including the cycling of organic materials such as carbon through decomposition of recalcitrant plant materials (e.g., cellulose, lignin, xylan). Bacteria and fungi associated with the top layer of soil – i.e., the leaf litter microbiome – break down plant saccharides, releasing carbon dioxide through respiration, which influences land-atmosphere carbon exchange (1). Therefore, studying saccharide degradation is key for predicting climate change feedbacks and the rate of carbon remineralization (2, 3).

During leaf litter decomposition, microbes incrementally decompose plant-derived saccharides and generate a shared pool of saccharides varying in substrate complexity. On the one end of the substrate complexity gradient are complex saccharides, defined as polymeric saccharides requiring external processing for microbial consumption (4). An example of complex saccharides is xylan, the second most abundant plant cell wall polysaccharide after cellulose. Xylan can vary in structural complexity from a linear chain of β-1,4-linked xylose (homoxylan) to a decorated xylose backbone with branching sugar residues other than xylose (heteroxylan). Xylans are degraded by carbohydrate-active enzymes (CAZymes), which hydrolyze specific bonds within the polysaccharide chain. For example, endo-1,4,-β-xylanase hydrolyzes the xylan backbone and converts xylan into an accessible form of xylose. On the other end of the substrate complexity gradient are simple saccharides, including the monomeric forms of the polysaccharides (e.g. xylose), that are easily transported inside the cell without external processing. While a broad range of species have the transporters to import mono- and oligosaccharides, fewer species produce CAZymes that degrade polysaccharides (here referred to as ’degraders’). Furthermore, degraders vary in the composition and abundance of their CAZymes, with the copy and type of genes varying widely among species.

Given that the degradation of complex saccharides involves CAZymes that act as public goods and any microbe in the vicinity can directly profit from the hydrolyzed products, the process of polysaccharide degradation has the potential to shape microbial behaviors and interactions (5, 6). For example, *Caulobacter crescentus* displayed different behaviors depending on its growth on the polysaccharide xylan or the monosaccharide xylose (5). In xylan, cells interact positively by co-aggregating to localize extracellular enzymes that would otherwise be lost to diffusion. In contrast, in xylose, cells interact negatively and disperse to avoid resource competition (5). Thus, substrate complexity influences interactions among closely related individuals.

Yet, its effect on interspecific interactions remains less understood. It is well-established that 1) simple carbohydrates promote negative interspecies interactions through resource competition (7) and 2) complex carbohydrates promote positive interactions between specialized degraders and downstream consumers, the latter of which lack the CAZymes necessary to degrade polysaccharides and, subsequently, cross-feed on simple saccharides released by the degraders (6). Beyond these observations, it is unknown how substrate complexity influences interspecific interactions among degraders. Here, we hypothesized three outcomes (Figure 1). In the no interactions hypothesis, degraders differentially hydrolyze and consume variable breakdown products from the polysaccharide that other degraders do not use (i.e. resource partitioning; Figure 1, H2a). Alternatively, positive interactions among degraders are expected when synergisms emerge from the division of labor between their CAZymes (4). For example, some CAZymes may initially degrade the branching sugar residues, facilitating other CAZymes to target other oligosaccharides (Figure 1, H2b). Finally, a combination of interactions is plausible (Figure 1, H2c).

**Figure 1.**
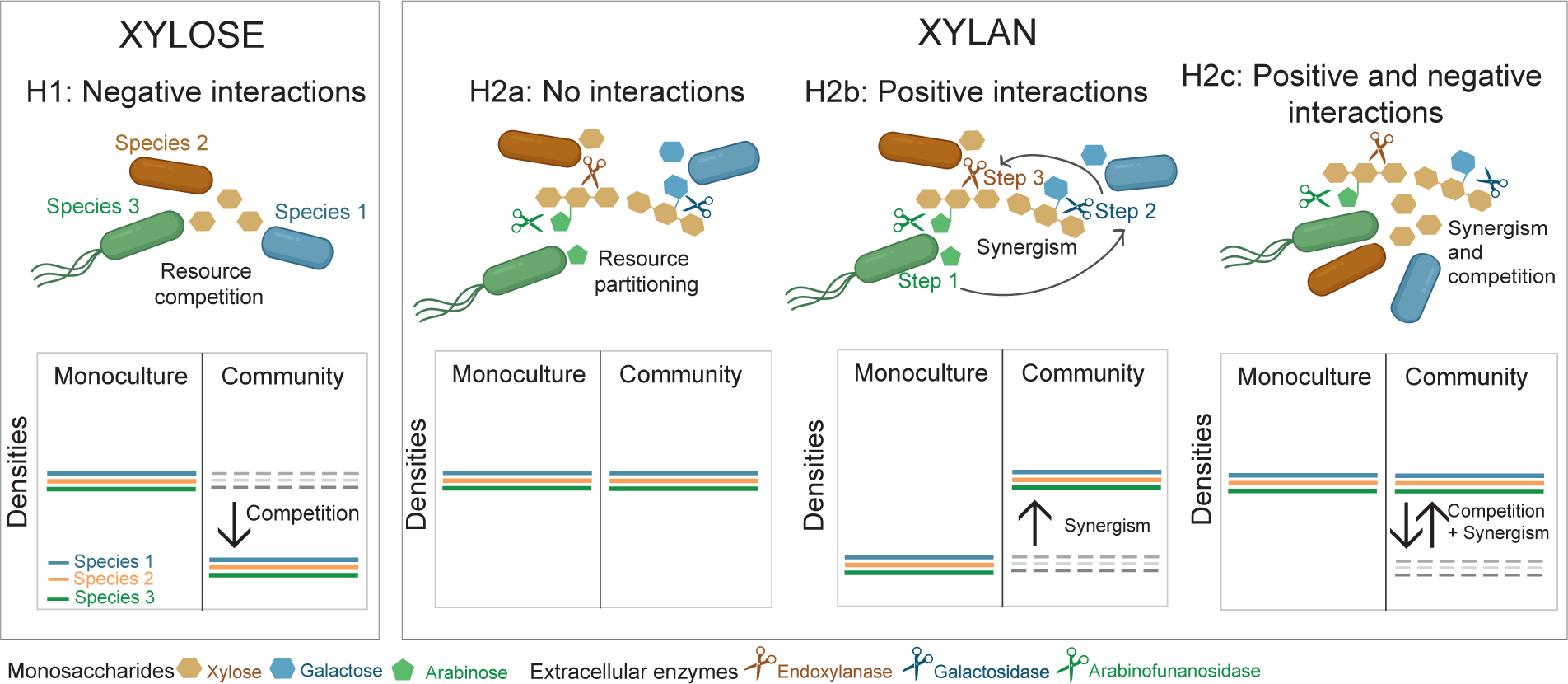
Expected interspecific interactions on simple and complex carbohydrates. In xylose, negative interactions among species are expected due to increased resource competition for simpler substrates (H1). Under this scenario, species’ densities in a community (represented by solid-colored lines) are expected to be lower than observed in monoculture. In xylan, several interactions may occur. First, no interactions are expected if species specialize in hydrolyzing and consuming part of the polysaccharide through differential production of various extracellular enzymes (represented by scissors) that differ in their hydrolytic enzyme activity. (H2a). Under this scenario, species’ densities are expected to be similar in the community than in monoculture. Second, positive interactions are expected if species cooperate to degrade the polysaccharide. Synergistic interactions can occur if the polysaccharide is degraded sequentially (H2b). Specialized CAZymes hydrolyze parts of the polysaccharide, and only after the hydrolysis products are removed the next hydrolysis step can proceed. Due to this synergism, species densities in the community are expected to be higher than in monoculture because species on their own cannot fully hydrolyze the polysaccharide. Finally, positive and negative interactions are expected when species act synergistically to degrade the polymer but compete for the breakdown products released (H2c). Under this scenario, positive and negative effects may cancel each other out so that species’ densities are similar in the community than in monoculture.

Understanding species interactions during polysaccharide degradation can also provide insights into how nutrient complexity affects patterns of species coexistence. Beyond simple theory, where an environment with only one resource can sustain one species due to competition (8), it is harder to predict coexistence patterns in polysaccharides. One could expect that polysaccharides promote species coexistence through niche partitioning (7), or because species’ growth is limited due to increased energetic investment in extracellular enzyme production to degrade polysaccharides. These expectations have rarely been directly tested.

Here, we study the influence of carbohydrate complexity on the interactions and coexistence among three xylan-degraders isolated from the leaf litter microbiome of a Mediterranean grassland site in Southern California (9). The leaf litter microbiome has been used as a model system to study microbial responses to climate change as it performs an important ecosystem function – the decomposition of plant polysaccharides – and because it is less complex compared to bulk soil (10, 11). Despite its relative simplicity, studying species interactions within soil communities is challenging (12). Therefore, we simplified the leaf litter microbiome by assembling a three-species synthetic community that is easy to manipulate and trackable. The three species – *Curtobacterium* sp., *Flavobacterium* sp., and *Erwinia* sp. – were selected for two reasons. First, the species belong to the most diverse and abundant phyla (Actinobacteria, Bacteroidetes, and Proteobacteria, respectively) from the grass litter communities comprising 95% of total bacteria abundance (13–15). At the genus level, *Curtobacterium* is the most abundant in the grassland and represents 8-12% of the bacterial cells on grass litter (14, 15). Second, the three species capture the metabolic diversity of the leaf litter microbiome. While most of the species from the leaf litter microbiome degrade complex polymers, species within each phylum share unique carbon use patterns (16). For instance, species from the genus *Curtobacterium* are excellent ‘degraders’ and breakdown complex polymers such as cellulose, hemicellulose (xylan), and other plant polysaccharides (17–19); whereas, γ-Proteobacterium species (e.g., *Erwinia*) are considered ‘consumers’ defined as fast growers that specialize on oligosaccharides and monosaccharides such as glucose and xylose (16). Finally, Bacteroidetes (e.g., *Flavobacterium*) are considered ‘generalists’ having the genomic potential to use a broad set of polysaccharides and oligosaccharides (19).

Using this simplified system, we showed that substrate complexity influenced species growth, mediated interspecific interactions, and promoted species coexistence in a synthetic model community that is relevant in the context of climate change.

## Materials and Methods

### Bacterial isolates

To assemble a simple community representative of the grass litter microbiome, we followed a bottom-up approach and selected three bacterial strains isolated from the Loma Ridge site (Table 1). The grassland site from the Loma Ridge Global Change Experiment (Irvine, California, USA, 33° 44’ N, 117° 42’ W), is dominated by the annual grasses from the genera *Avena*, *Bromus*, and *Lolium*, and the perennial grass *Nassella pulchra* (9, 16). Strains were isolated from surface litter as previously described (16) and stored at -80°C in glycerol stocks (32.5%) for genomic and culturing analyses.

**Table 1.** Bacterial isolates used in this study.

| Phylum<br>(Average relative abundance of Phylum) | Family | Genus<br>(Relative Abundance of Genus) | Species<br>(strain) | Accession number <sup>4</sup><br>(identifier) | Previous studies that have characterized the isolate |
| --- | --- | --- | --- | --- | --- |
| Actinobacteria<br>(48.0% <sup>1</sup> – 51.5 % <sup>2</sup>) | Microbacteriaceae | *Curtobacterium*<br>(10.4% <sup>3</sup> - 18.64% <sup>2</sup>) | *Curtobacterium* sp.<br>(MMLR14002) | PRJNA524191<br>MMLR14_002<br>NZ_MJGT000000000 | Ref. 15, 17, 18 and 20 |
| Bacteroidetes<br>(6.8% <sup>1</sup> – 26.5 % <sup>2</sup>) | Flavobacteriaceae | *Flavobacterium*<br>(0.78 % <sup>2</sup> - 5.5 % <sup>3</sup>) | *Flavobacterium* sp.<br>(LR40) | PRJNA391502<br>MMLR14_040<br>SAMN38146147 |  |
| Proteobacteria<br>(25% <sup>2</sup> – 35.4% <sup>1</sup>) | Erwiniaceae | *Erwinia*<br>(<0.01 % <sup>2</sup>) | *Erwinia* sp.<br>(LR17) | PRJNA391502<br>MMLR14_017<br>SAMN38146146 | Ref. 16 |

<sup>1</sup> Average from metagenomic sequencing data (15)
<sup>2</sup> From natural litter pyrosequencing data (14)
<sup>3</sup> From 16S sequencing of the grassland microbial community. https://doi.org/10.1525/elementa.2021.00124
<sup>4</sup> Gene Bank accession number: BioProject, Organism, Accession number (this and previous studies)

The genome of *Curtobacterium* MMLR14002 was previously sequenced (GeneBank’s accession NZ_MJGT00000000, ref. 20). We sequenced the genomes of *Flavobacterium* sp. and *Erwinia* sp. (accession numbers SAMN38146147 and SAMN38146146, respectively). Bacterial isolates were revived from glycerol stocks and streaked on Lysogeny Broth (LB) agar plates. A colony was grown in 10 ml of LB media for 48 h at 26°C with shaking (100 r.p.m.). We used 1 ml of cultures and isolated the genomic DNA using the Wizard Genomic DNA Purification Kit (Promega; Madison, WI). We used the DNA with the Illumina DNA Prep kit (Illumina; San Diego, CA, USA) with the low-volume protocol (dx.doi.org/10.17504/protocols.io.bvv8n69w) for library preparation that was quality assessed using a Bioanalyzer, prior to sequencing on the NovaSeq6000 S4 flow cell.

Raw reads were processed for quality control (trimq=32 qtrim=rl) and adapter removal (ktrim=r k=25) with bbduk.sh (21) and used as input for de novo assembly with SPAdes (22) by increasing k-mers from 31-111 under the “careful” h-path removal strategy to reduce mismatches and indels. Draft assemblies were screened for quality by constructing taxon-annotated GC-coverage plots using blastn (23) and bowtie2 (24), respectively. Based on the resulting metrics, we removed all contigs <5000bp and coverage <30 for further analyses. Genome completeness and contamination were calculated checkM (25) and taxonomy was verified using the gtdbtk classifier (26). To maintain consistency across genomes, all genomes were assigned open reading frames (ORFs) using prodigal (27) and gene annotations with prokka (28).

### Whole genome analysis

To identify strains’ genomic potential for carbohydrate degradation, we screened all ORFs against the protein family database, Pfam-A (29), using hidden Markov models, hmmsearch (30). Resulting protein domains were filtered for their top match based on e-value and bitscore. Using a curated list, we queried the top domain hits for the presence of glycoside hydrolase (GH) and carbohydrate binding module (CBM) domains in each strain (18). This approach identified GH/CBM families with their associated enzymatic activity and target substrate (31). Potential xylan degraders were defined as bacteria having at least one GH/CBM gene targeting xylan (Table S2).

### Growth conditions

Strains were grown in a modified M63 medium supplemented with either xylan or xylose. The xylan from the corn core used in this study was composed of 76% xylose after hydrolysis (Tokyo Chemical Industry). The chemical composition of this modified M63 medium was: 47 mM K_2_HPO_4_, 39 mM KH_2_PO_4_, 15 mM (NH_4_)_2_SO_4_, 1.8 µM FeSO_4_, 1 mM MgSO_4_, 34 nM CaCl_2_, 92 nM Fe III Chloride, 0.73 nM ZnCl_2_, 6.3 nM CuSO_4_ 5H_2_0, 8.4 nM CoCl_2_ 6H_2_0, 4.1 nM Na_2_MoO_4_ 2H_2_0, 5.9 µM thiamine, and 0.02 % peptone. Peptone was added to the medium as a source of amino acids. All solutions were sterilized using 0.22 µm PES filters. Batch culture experiments were performed in 40 ml glass vials with screw caps containing TFE-lined silicone septa (32).

### Growth curves

We characterized bacterial growth at different concentrations of xylan and xylose (from 0 to 10 mM). We revived the bacteria by streaking 10 µl of glycerol stock onto LB agar plates and incubating the plates at room temperature. We then picked a colony from each species and grew them in 10 ml of fresh M63 medium with 4 mM xylose. Cultures were incubated for 48 hours at 26°C with shaking (100 r.p.m.). We transferred 2 µl of the saturated culture into 198 µl of fresh media dispensed in each of the wells of the 96-wells plate. To avoid evaporation, we covered the plates with a lid, which we sealed with silicon grease. We incubated the plates at 30°C with shaking inside a photospectrometer plate reader (Epoch2^TM^, Agilent^TM^). Optical Density (OD_600_) measurements were acquired every 10 minutes for 72 hours.

From each growth curves, we estimated: 1) the maximum growth rate *µ_max_*, defined as the maximum growth rate during the exponential growth phase (h^-1^); 2) the maximum population size *OD_max_*, defined as the maximum OD after saturation minus the minimum OD at the start of the experiment (unitless OD); 3) the yield *Y*, defined as the amount of biomass produced per unit of resource (unitless OD/mM). Three growth curves (technical replicates) were used to get an average of each parameter.

### Co-culture invasion experiments

To determine if species coexisted in pairs, we did co-culture invasion experiments in batch culture. We co-inoculated species at varying initial fractions so each species was underrepresented, equally represented, and overrepresented. To reach these initial fractions, we first determined the relationship between the OD and the cell density estimated by colony forming units (CFUs) counts. To do so, we grew each species in M63 medium with 16.67 mM of xylose to reach high bacterial densities. We then used the saturated cultures and diluted them to obtain two-, four-, and eightfold dilutions. We measured the OD from these dilutions in the plate reader and plated them in LB agar to estimate their CFU. We replicated the experiment two times and fitted a linear relationship between the OD and the cell density for each species (Figure S1).

To start the invasion experiments, we acclimated cultures by inoculating a colony from each species into 10 ml fresh M63 medium with 16.67 mM xylose and incubated them for 48 hours. We used the OD/CFU relationship (Figure S1) to calculate the volumetric ratios to reach the targeted initial ratios of 1:10, 1:1 and 10:1. The co-cultures were then incubated at 26°C with shaking (100 r.p.m.). After 48 h, we transferred 0.1% of the population into a fresh medium and repeated this transfer again to obtain three growth-dilution cycles (i.e., six days). After each cycle, we estimated the species densities by plating cultures on LB plates and determining the CFU for each species. This was possible given that each species has a unique colony size and morphology on LB plates (Figure S2). Species densities were used to estimate relative abundances for pairs of species estimated as fractions; e.g., fraction species A = cell density species A / (cell density species A + cell density species B). Initial fractions were 0.1 ± 0.1 (rare), 0.5 ± 0.1 (equal), and 0.9 ± 0.1 (common). An experimental duplicate for each of the three initial fractions was obtained. Changes in relative abundances over time were used to determine the type of competition outcome (33). Stable coexistence occurred when one species increased in frequency when rare and decreased in frequency when common until it reached a coexistence equilibrium significantly different from 0 or 1 (one-tailed t-test). Competitive exclusion was observed when one species increased in frequency regardless of its initial frequency and reached a final fraction non-different from 1 (one-tailed t-test). Statistical analyses were done using R (version 4.1.3).

### Pairwise species interactions

We quantified ecological interactions between pairs of species by comparing the densities of species grown alone (monocultures) or with a second species (pairwise cocultures). Cultures were grown in batch culture as described in the previous section. We compared the densities (CFU/ml) at day 6 in monoculture and co-culture, performing a t-test over the means (three replicates). If the densities were not significantly different in both monocultures and co-cultures, we concluded that species had no interaction. If the densities in co-cultures were significantly lower than in monocultures, we concluded that species had a negative interaction.

### Three-species community experiments

To determine if all three species coexisted in a community, we did batch culture experiments over 10 days. We also determined the effect that other species had on growth by comparing species densities in monoculture and in a community. Monocultures were started by diluting the acclimated cultures 1000-fold into 10 ml M63 medium with xylose or xylan. In communities, the three species were mixed at an equal ratio by adjusting the volumes of each species to reach an initial target population density of 10^5^ cells/ml per species. Cultures were incubated at 26°C with shaking (100 r.p.m.) for 48 hours. Monocultures and communities were propagated by transferring 0.01 mL of the culture into 9.99 mL of fresh medium every 48 hours for a total of 10 days. We repeated these experiments two more times to obtain three temporal replicates.

To define the effect that other species have on growth, we compared the densities over 10 days for each species in monoculture and in the community and performed a two-tailed t-test over the means. If the densities were not significantly different between a species grown in monoculture and in the community, we concluded that species in the community had no effect on the species of interest. If the densities in monocultures were higher than in communities, we concluded that species in the community had a negative effect on the species of interest. Statistical analyses were done using R (version 4.1.3).

### Resource explicit model

We built a mathematical model that explicitly modeled the changes in resources and the growth of the three species. We parameterized the model using single-species growth measurements derived from growth curves in xylose (4mM) and xylan (4mM). We estimated three parameters for each of the species: 1) the maximum growth rate (*µ_max_*), 2) the maximum uptake rate (*V_max_*) the half-saturation constant (*k*). Each parameter was fitted to three growth curves (i.e. technical replicates) as (32) using Matlab (version R2017a, Mathworks), except for *µ_max_*. *µ_max_* was estimated by first calculating *µ*, the instantaneous growth rate (h^-1^), as Δ ln(OD)/Δ*t* with Δ*t* = 0.16 h^-1^ (10 min) and then selecting the maximum value of *µ* in the exponential phase.

In our mathematical model, the growth of each species was dependent on the concentration of resources, which was modeled explicitly using Matlab’s ODE45 solver. Bacteria and resource concentrations were modeled with the following differential equations (Table S1). We incorporated the daily dilution that resulted from transferring 1/1000 of the culture into fresh media every 48 hours and added an extinction threshold in which extinction occurred when less than one bacterium was transferred to the next cycle. Finally, to compare the results generated by the mathematical model to the experimental data, we converted the numerical values generated by the model (in OD units) to cell densities (in CFU) using the slope of the linear relationship between the OD and the cell density as the conversion rate (Figure S1).

## Results

### Species in monoculture are capable of degrading xylan but utilize it less efficiently than xylose

First, we assessed the genomic potential for xylan degradation of the three isolates by characterizing the abundance and composition of carbohydrate binding module (CBM) and glycoside hydrolase (GH) protein families associated with xylanase activity. The abundance of CBM/GH genes involved in xylan degradation varied among genomes, with *Flavobacterium* encoding the most xylanases (34 CBM/GH xylanases copies), followed by *Curtobacterium* (6 CBM/GH xylanases copies) and *Erwinia* (2 CBM/GH xylanases copies; Table S2). All strains had at least two copies of endo-1-4-xylanases ("true xylanases"), which degrade the xylan backbone (34). Based on the presence of xylanase genes, all three species were classified as potential xylan degraders.

To determine if genomic potential translated into degradation, we grew the species in monoculture with different concentrations of xylan (Figure S3). The maximum population size increased with the concentration of xylan supplied, indicating that xylan was a limiting growth factor in our monocultures (Figure S3). While all three species utilized and degraded xylan as their main source of carbon and energy (35), there was high variability in growth among species. For example, at 4 mM of xylan, the maximum growth rate and population size significantly varied among species (one-way ANOVA, p=2.0 x 10^-4^ and p=4.6 x 10^-12^, respectively). *Curtobacterium* and *Flavobacterium* were efficient xylan degraders and achieved a higher maximum population size than *Erwinia* (Figure 2; Figure S3). Unexpectedly and unlike the other strains, *Curtobacterium* also grew in the control treatment without xylan (Figure S3). That is, *Curtobacterium* utilized peptone as a carbon source. *Curtobacterium*’s ability to use peptone as a source of carbon resulted in a diauxic shift when grown in xylan and peptone (Figure 2, Figure S3), with the growth on peptone accounting for approximately half of its total growth. Therefore, after we normalized *Curtobacterium* growth by accounting for peptone, *Curtobacterium* and *Flavobacterium* achieved similar maximum population sizes. Still, *Curtobacterium* had a higher maximum growth rate than *Flavobacterium* (Figure S3). In sum, even though all three species utilized xylan – i.e. they are all xylan degraders – they each had unique growth behaviors. *Curtobacterium* grew fast and achieved a high final population size, *Flavobacterium* grew slow and achieved a high final population size, and *Erwinia* grew moderately fast and achieved a low final population size.

**Figure 2.**
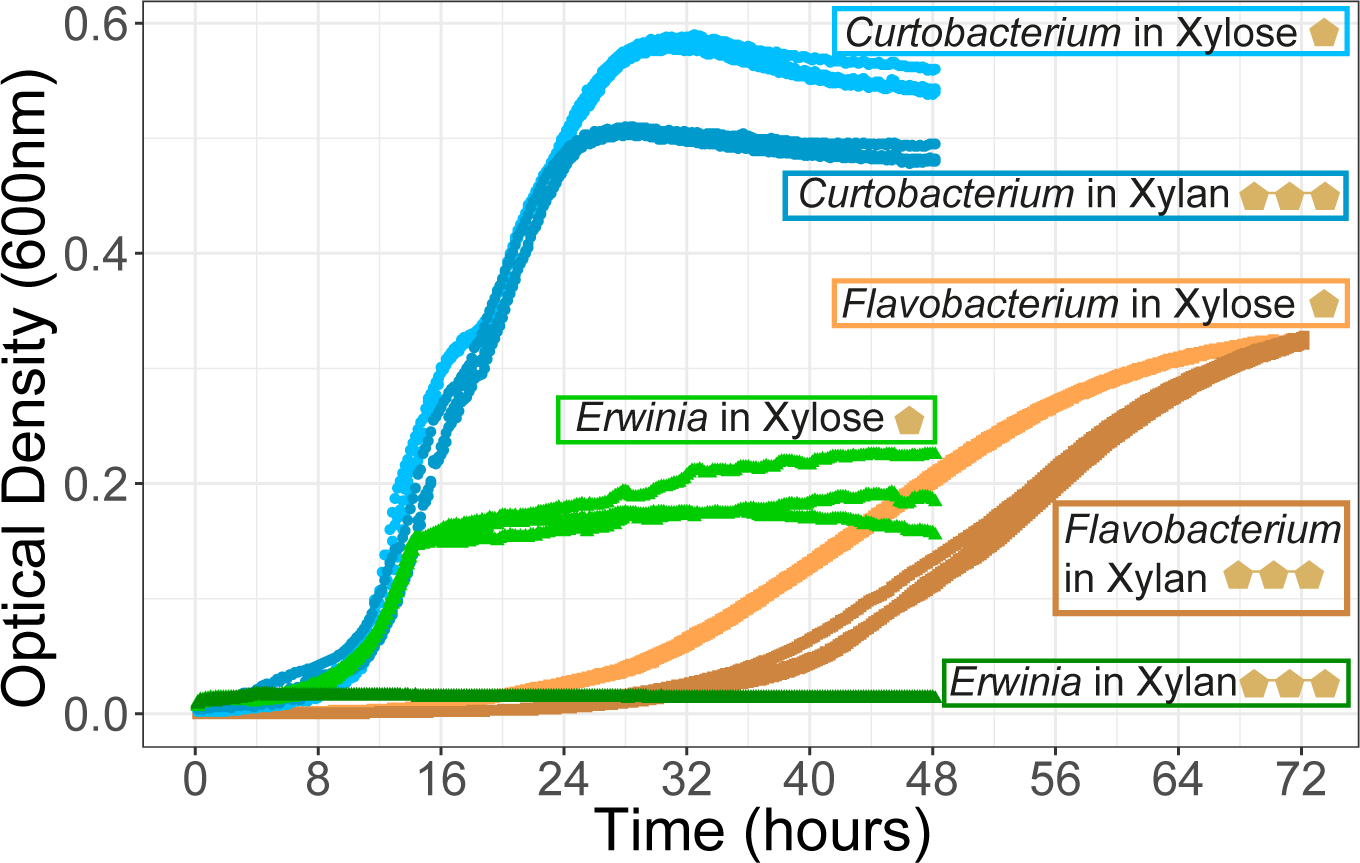
Species have unique growth profiles in xylan and xylose. Growth curves of species grown in isolation on 4 mM of the polymer xylan and on 4 mM of its constituent monomer xylose. A close-up of *Erwinia*’s growth on xylan is available in Figure S3. The optical density measured at 600 nm over 48 hours is plotted for three technical replicates per species.

Next, we grew the species in monomer xylose at the same concentrations (weight/volume) previously used for the polymer xylan. Overall, the three species grew better in xylose than in xylan, even if both carbon sources were provided at the same concentration (Figure S3). *Erwinia* sp. achieved a 22-fold higher maximum population size in 4 mM xylose than in 4 mM xylan (0.136 unitless OD in xylose; 0.006 unitless OD in xylan; two-tailed t-test, d.f. = 2, and *P* = 3.063 x 10^-5^; Figure 2). The growth limitation in xylan was less severe but significant for *Curtobacterium* with a 1.2-fold lower maximum population size in 4 mM xylan than in 4 mM xylose (two-tailed t-test, d.f. =2, *P* = 1.847 x 10^-5^; Figure 2). *Curtobacterium* and *Erwinia*, also grew significantly slower in 4mM xylan than in 4mM xylose (two-tailed t-test on the maximum growth rate, d.f. =2, *P* = 0.001 and *P* = 0.003, respectively). In turn, *Flavobacterium* had a higher maximum growth rate in xylan than in xylose (Figure S3), but it had a longer lag phase in xylan than in xylose (Figure 2).

### All pairs of species stably coexist in xylan, but two pairs outcompete each other in xylose

Next, we assessed if species could coexist when grown in co-culture in xylan or xylose. We used 4 mM of xylan and xylose, which is a concentration below the saturation threshold for all three species (Figure S3). All pairs stably coexisted in xylan (Figure 3). For each pair, both species invaded each other when starting at low initial frequencies. On day six, species converged into a final fraction regardless of the starting fractions. For all pairs, the final fraction was skewed towards the dominance of one species. For example, *Curtobacterium* was overrepresented when growing in xylan and reached a final fraction of 0.93 ± 0.02 in pair with *Erwinia* and 0.82 ± 0.02 in pair with *Flavobacterium* (Figure S6). For the *Flavobacterium*-*Erwinia* pair, the coexistence equilibrium was shifted towards *Flavobacterium* being overrepresented (final fraction of 0.87 ± 0.03). *Erwinia* was always underrepresented (Figure 3). In contrast to xylan, coexistence patterns differed in xylose, with two pairs displaying competitive exclusion. In both cases, *Flavobacterium* was outcompeted. Only *Curtobacterium* and *Erwinia* stably coexisted in xylose (Figure 3).

**Figure 3.**
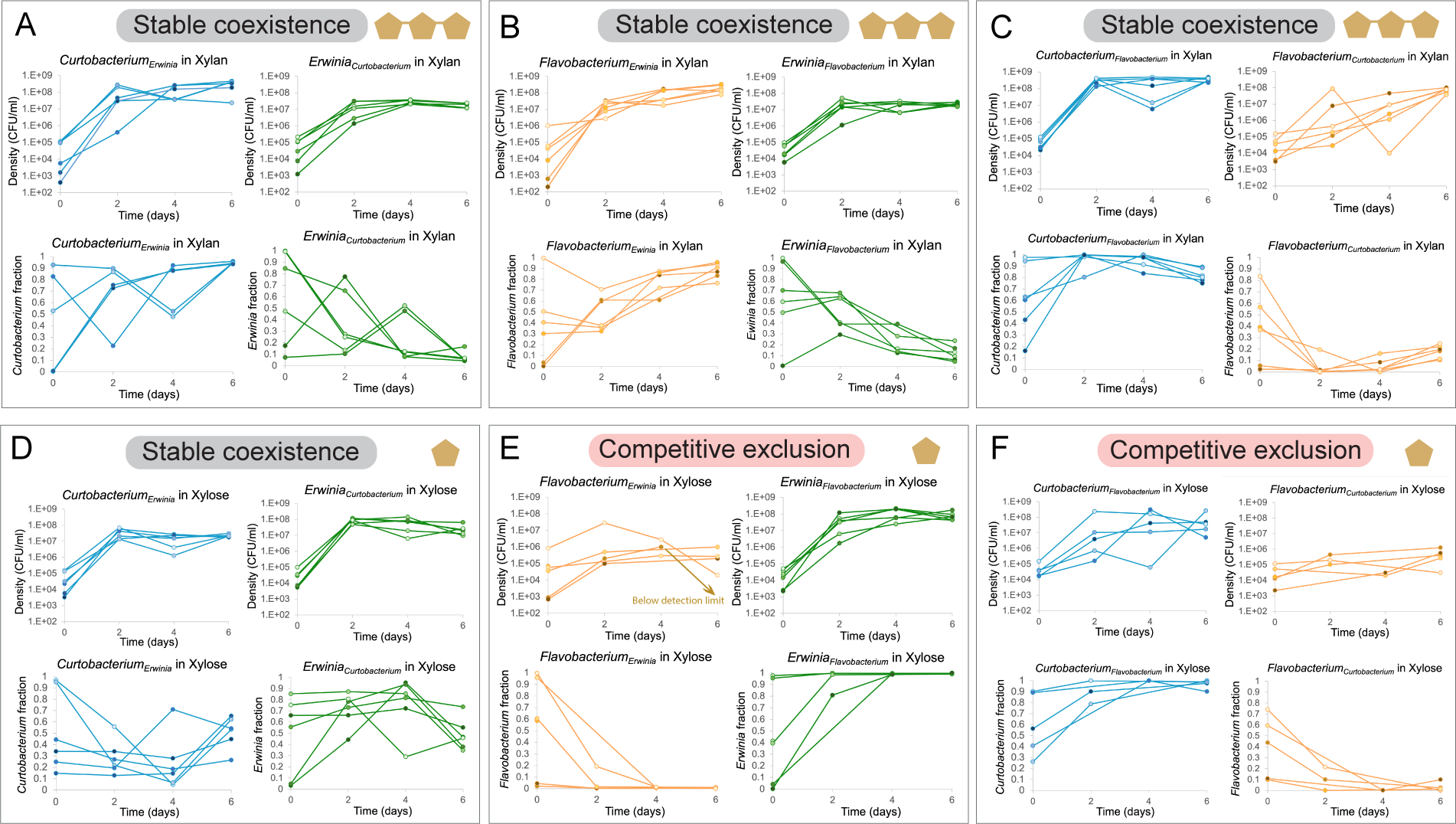
Pairwise competitions in xylan result in stable coexistence whereas, in xylose, they result in competitive exclusion or stable coexistence. Pairwise competitions in xylan and xylose, respectively, for the *Curtobacterium*-*Erwinia* pair (panels A and D), *Flavobacterium*-*Erwinia* pair (panels B and E), and *Curtobacterium*-*Flavobacterium* pair (panels C and F). For each panel, the upper graphs represent the population density (CFU ml^-1^) trajectories of species in co-culture over 6 days for six replicates. The lower graphs represent the fraction (values between 0 and 1) of each competing species over 6 days calculated from the population densities (upper panel). In stable coexistence, a species’ fraction increases when started from rare and decreases when started from common until an equilibrium fraction significantly different from 0 or 1 is reached (panels A to D). In competitive exclusion, the fraction of the strongest competitor increases until it reaches a fraction of 1 regardless of whether it started from rare or common (panels E and F). Competitive exclusion results in the extinction of *Flavobacterium* for one of the replicates in the *Flavobacterium*-*Erwinia* pair (panel E).

### Pairwise interactions are negative in xylose and not significant in xylan

We further characterized the network of pairwise interactions by comparing the growth of species in monoculture to its growth in pairwise co-culture. For all pairs in xylan, there were no significant differences in the final densities in monoculture and co-culture (Figure 4 A). In contrast, in xylose, negative pairwise interactions prevailed. *Erwinia* had significantly lower densities in co-culture with *Curtobacterium* than when growing in monoculture (2.7-fold lower in the co-culture than in monoculture, two-tailed t, d.f. 8, *P* = 0.003). *Flavobacterium* also reached lower densities in co-culture with *Erwinia* and *Curtobacterium* than in monoculture, but these differences were marginally significant (Figure 4 B). This lack of significance is likely due to a large variability among replicates, including some replicates for which the final densities dropped below the detection limit. Neither *Erwinia* nor *Curtobacterium* were affected by the presence of other species. Thus, in xylose, interactions between pairs of species were amensalistic.

**Figure 4.**
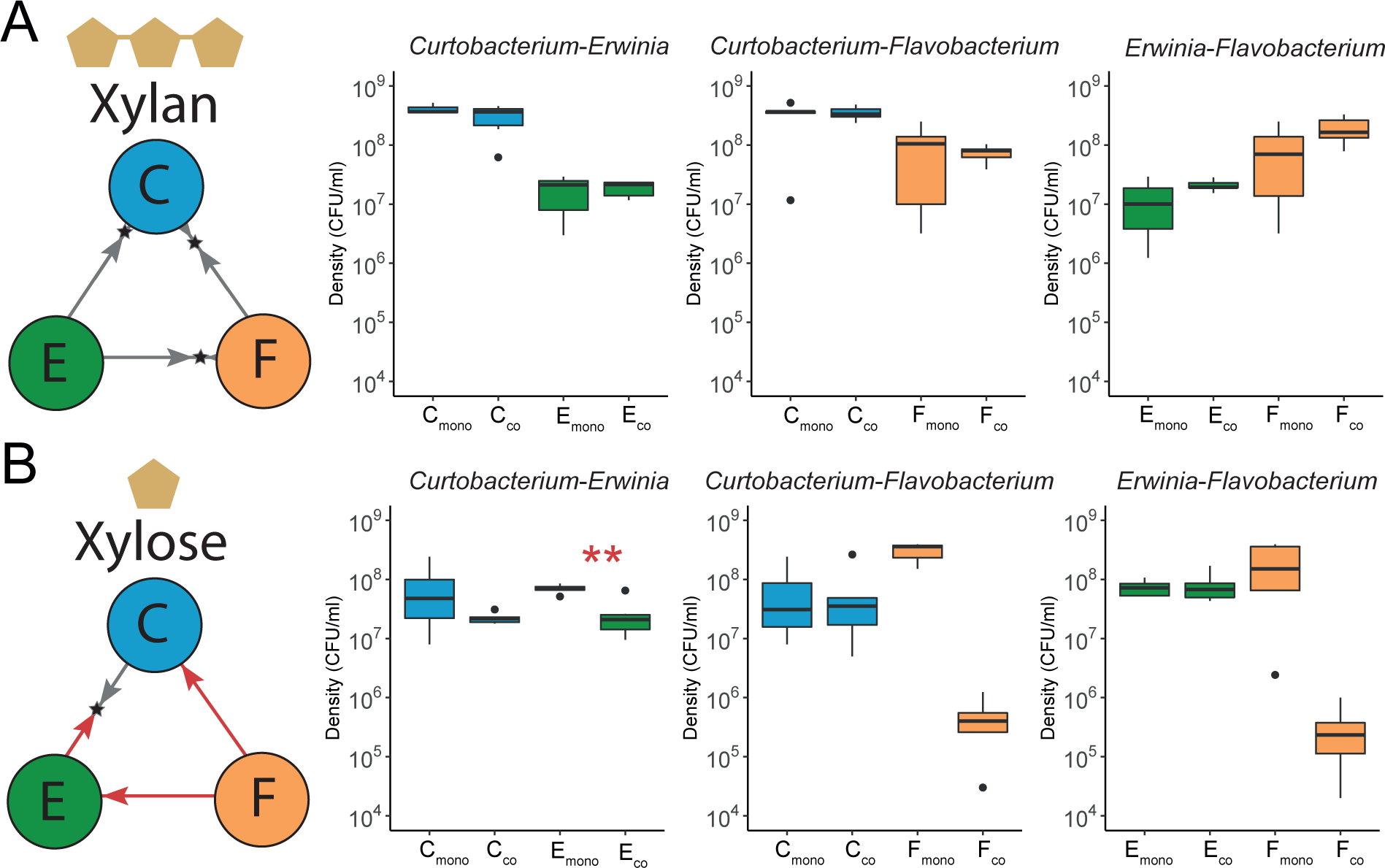
The pairwise interaction network differs in xylan and xylose. In xylan (panel A), pairwise competitions result in stable coexistence of *Curtobacterium* (C), *Erwinia* (E), and *Flavobacterium* (F), which is represented by two arrows pointing to a star representing the final fraction on day six (summary from Figure 3). The growth in monoculture and in co-culture at day six is represented by box plots (average of three to six replicates). For all three species, the growth in monoculture (C_mono_, E_mono_ and F_mono_) is not statistically different than the growth in co-culture (C_co_, E_co_ and F_co_). Thus, interactions between species were not significant in xylan. Instead, in xylose (panel B), pairs interact negatively. *Flavobacterium* (F) was outcompeted by both *Curtobacterium* (C) and *Erwinia* (E), which is represented by a single red arrow (summary from Figure 3). The growth of *Erwinia* at day six in co-culture with *Curtobacterium* was significantly lower than its growth in monoculture (*P* = 0.003; Welch Two Sample t-test). The growth of *Flavobacteriu*m at day six in co-culture with the other species was lower, but these differences were only marginally significant (*P* = 0.068 and *P* = 0.058; Welch Two Sample t-tests).

### All three species coexist in xylan, whereas only two species coexist in xylose

Next, we assembled a three-species community and compared species’ growth in the community and in monoculture. In xylan, the three species grown in a community sustained a high and stable population over 10 days (Figure 5A, right panel). These population densities did not differ to those in monocultures (two-tailed t-test, *P* = 0.831, *P* = 0.316 and *P* = 0.123, respectively), suggesting that other species had no effect on a species’ net growth. In contrast, in xylose, growth was diminished in the three-species community compared to monocultures (Figure 5B). *Curtobacterium* sustained significantly lower densities in the community than in monoculture (8-fold lower growth in the community than in monoculture, two-tailed t-test, d.f. 14, *P* = 0.022). *Erwinia* also had a lower density in the community than in monoculture (1.6-fold lower growth in the community than in monoculture), even though the differences were not statistically significant. Finally, *Flavobacterium* reached a density 50 times lower in the community than in monoculture (two-tailed t-test, d.f. 14, *P* = 0.011). *Flavobacterium* was the most impacted by the presence of other species, and its density dropped below the detection limit after the fourth serial transfer (Figure 5B, right panel). These negative effects of growing in a community were also captured by analyzing the area under the curve over ten days (Figure S4). In sum, all three species coexisted in xylan, whereas only two of them coexisted in xylose over ten days.

**Figure 5.**
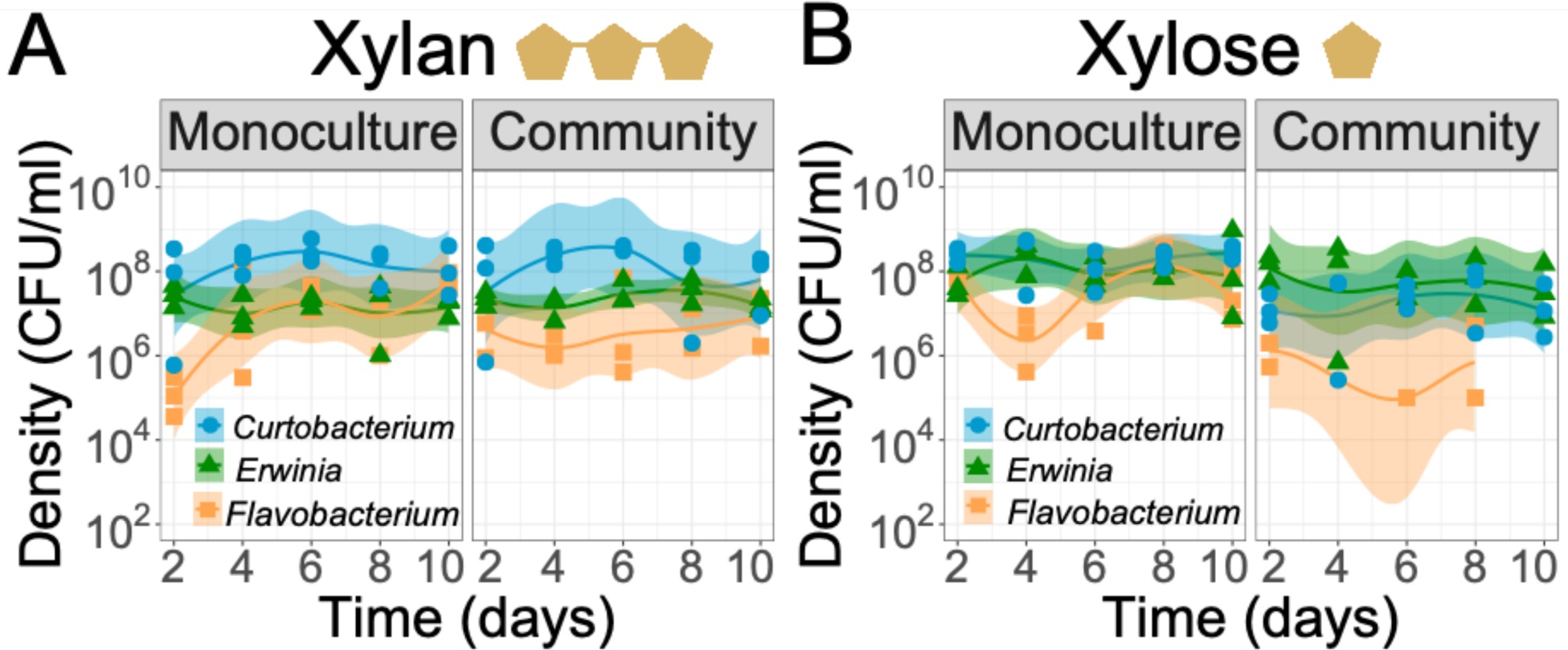
Coexistence patterns in a three-species community change according to the substrate complexity. In xylan, all three species coexisted and grew equally well in a three-species community than alone (panel A). Instead, in xylose, only two species coexisted for 10 days, and species grew less well in the community than alone (panel B). For each panel, population density trajectories of monocultures are represented on the left side and of communities on the right side, estimated from CFU mL^-1^ during five serial transfers (1000-fold dilution every two days). Each line corresponds to a local polynomial regression fitting of three replicates with 95% confidence interval. Graphs were constructed with the function `stat_smooth` (R version 3.2.3).

### A mathematical model recapitulated the community dynamics observed experimentally in xylose but not in xylan

To provide a mechanistic understanding of species interactions in communities, we built a null mathematical model to determine if resource competition alone can recapitulate our observed community dynamics. We simulated a simple case of resource competition in which the resources are readily used, regardless of whether they are in polymeric or monomeric form. Our model does not account for the production and release of costly extracellular enzymes for xylan degradation, nor the pool of shared xylose derived from the activity of these ’public goods’. With this null model we can assess if the better growth of monocultures in xylose vs xylan (Figure 2) suffices to explain why only two species coexist in xylose compared to xylan (Figure 5). Any deviation from our simple model would suggest our system is more nuanced and that other interactions besides resource competition may contribute to the observed community patterns.

We parameterized our mathematical model by using single-species growth behaviors in xylan and xylose (Table 2). When simulating five serial passages of communities in xylose, *Flavobacterium* went extinct and only *Curtobacterium* and *Erwinia* coexisted over ten days (Figure 6B). This result validate our experimental findings, except for the time of extinction of *Flavobacterium* which occurred after four days in the model (Figure 6B) compared to after eight days in the experiments (Figure 5B). However, in xylan, our model predicted the extinction of both *Flavobacterium* and *Erwinia* after four and six days, respectively (Figure 6A). This finding contrasts with our experimental data, where we observed coexistence of all three species over ten days (Figure 5A). This may be due, in part, to our model inaccurately reflecting monoculture dynamics; for example, *Flavobacterium* could not accurately be predicted in monoculture based on fitted parameters from our growth curves (Figure S5). However, *Erwinia*’s growth was accurately recapitulated in the monocultures (Figure S5), suggesting the persistence of *Erwinia* may be attributed to more complex interactions in xylan. Instead, in xylose, community dynamics could be mostly recapitulated with resource competition.

**Table 2.**
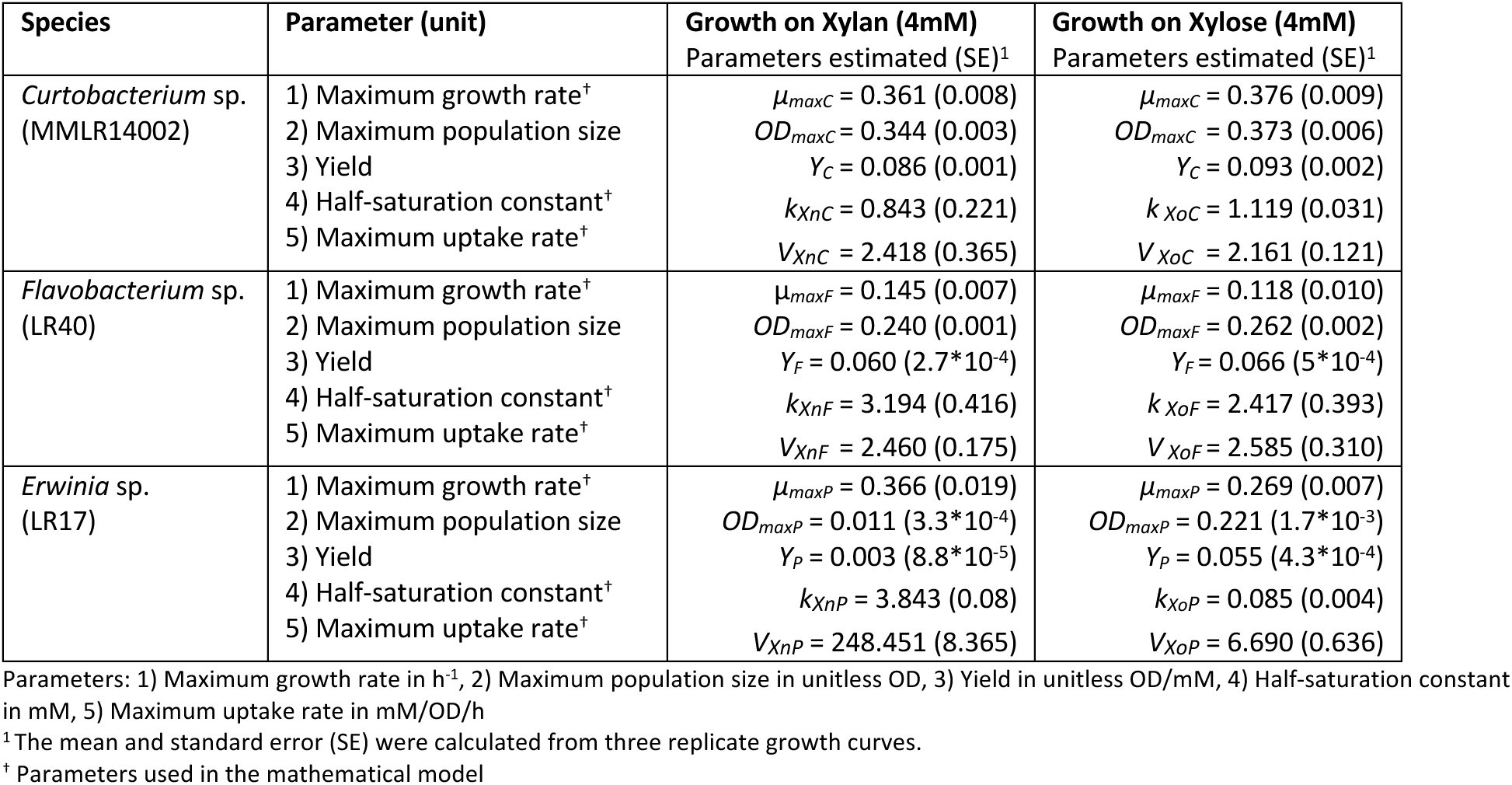
Growth parameters estimated from growth curves in xylan (4mM) and xylose (4mM).

| Species | Parameter (unit) | Growth on Xylan (4mM)<br>Parameters estimated (SE)<sup>1</sup> | Growth on Xylose (4mM)<br>Parameters estimated (SE)<sup>1</sup> |
| --- | --- | --- | --- |
| *Curtobacterium* sp.<br>(MMLR14002) | 1) Maximum growth rate<sup>†</sup><br>2) Maximum population size<br>3) Yield<br>4) Half-saturation constant<sup>†</sup><br>5) Maximum uptake rate<sup>†</sup> | $\mu_{maxC} = 0.361$ (0.008)<br>$OD_{maxC} = 0.344$ (0.003)<br>$Y_C = 0.086$ (0.001)<br>$k_{XnC} = 0.843$ (0.221)<br>$V_{XnC} = 2.418$ (0.365) | $\mu_{maxC} = 0.376$ (0.009)<br>$OD_{maxC} = 0.373$ (0.006)<br>$Y_C = 0.093$ (0.002)<br>$k_{XoC} = 1.119$ (0.031)<br>$V_{XoC} = 2.161$ (0.121) |
| *Flavobacterium* sp.<br>(LR40) | 1) Maximum growth rate<sup>†</sup><br>2) Maximum population size<br>3) Yield<br>4) Half-saturation constant<sup>†</sup><br>5) Maximum uptake rate<sup>†</sup> | $\mu_{maxF} = 0.145$ (0.007)<br>$OD_{maxF} = 0.240$ (0.001)<br>$Y_F = 0.060$ ($2.7 \cdot 10^{-4}$)<br>$k_{XnF} = 3.194$ (0.416)<br>$V_{XnF} = 2.460$ (0.175) | $\mu_{maxF} = 0.118$ (0.010)<br>$OD_{maxF} = 0.262$ (0.002)<br>$Y_F = 0.066$ ($5 \cdot 10^{-4}$)<br>$k_{XoF} = 2.417$ (0.393)<br>$V_{XoF} = 2.585$ (0.310) |
| *Erwinia* sp.<br>(LR17) | 1) Maximum growth rate<sup>†</sup><br>2) Maximum population size<br>3) Yield<br>4) Half-saturation constant<sup>†</sup><br>5) Maximum uptake rate<sup>†</sup> | $\mu_{maxP} = 0.366$ (0.019)<br>$OD_{maxP} = 0.011$ ($3.3 \cdot 10^{-4}$)<br>$Y_P = 0.003$ ($8.8 \cdot 10^{-5}$)<br>$k_{XnP} = 3.843$ (0.08)<br>$V_{XnP} = 248.451$ (8.365) | $\mu_{maxP} = 0.269$ (0.007)<br>$OD_{maxP} = 0.221$ ($1.7 \cdot 10^{-3}$)<br>$Y_P = 0.055$ ($4.3 \cdot 10^{-4}$)<br>$k_{XoP} = 0.085$ (0.004)<br>$V_{XoP} = 6.690$ (0.636) |

Parameters: 1) Maximum growth rate in h<sup>-1</sup>, 2) Maximum population size in unitless OD, 3) Yield in unitless OD/mM, 4) Half-saturation constant in mM, 5) Maximum uptake rate in mM/OD/h
<sup>1</sup>The mean and standard error (SE) were calculated from three replicate growth curves.
<sup>†</sup> Parameters used in the mathematical model

**Figure 6.**
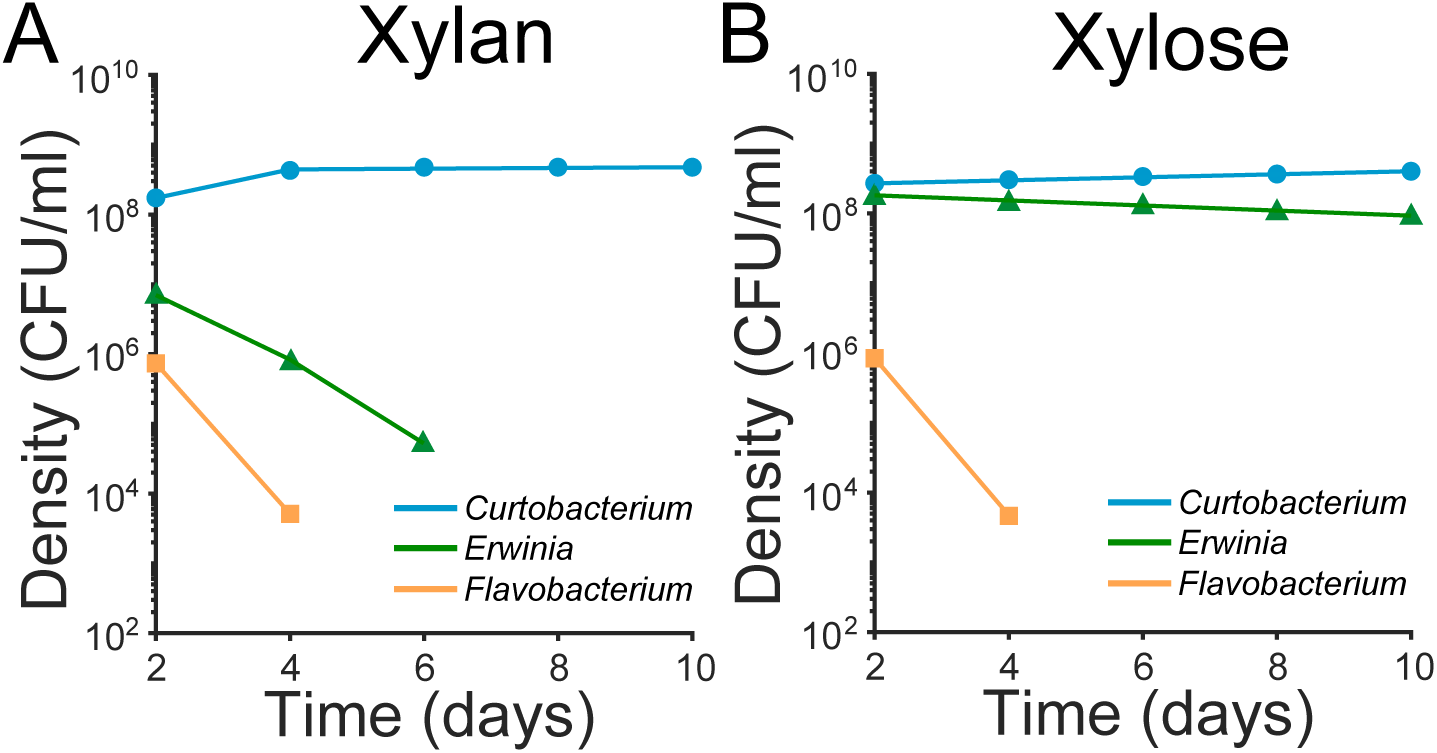
The mathematical model recapitulates community dynamics in xylose but not in xylan. The results derived from the mathematical model are plotted when simulating growth in a three-species community during five serial transfers in xylan (panel A) and in xylose (panel B). Each tick line corresponds to the connection between the final densities of each species after a 48-hour cycle of growth.

## Discussion

There is a growing interest in understanding how interspecies interactions shape the degradation of biopolymers in nature. Recent studies have focused on studying the interactions among multiple microbial guilds; for example, the interaction among degraders, exploiters and scavengers in marine polysaccharide-degrading bacteria (6, 36, 37). Here, we explored the interactions among three species belonging to the guild of xylan degraders from a natural leaf litter community that mediate the turnover of organic matter. We found that the complexity of the substrate influences the interactions and coexistence of species.

### Substrate complexity influences patterns of coexistence with less coexistence in xylose than in xylan

The complexity of the carbon source influenced the coexistence patterns: three species coexisted in xylan; instead, only two species coexisted in xylose. According to the coexistence theory, it is expected that, an environment with a single carbon source could only sustain one species. At first, it seemed surprising to see more coexistence than expected. Nevertheless, it is important to recognize that xylan/xylose may not act as the only carbon source, as *Curtobacterium* grew on peptone without an additional source of carbon. Thus, based on our system, we could expect that at least two species coexist, one of them being *Curtobacterium*. This is what we observed from experiments in pairs and communities in xylose. The observation that three species coexist in xylan (instead of two) could be explained either because xylan is a polymer or because the xylan in our study is composed of a chain of xylose decorated with branches of other carbon sources such as glucose and arabinose (38). Even if these other carbon sources are the minority (24%), it is possible that these extra carbon sources sustained the coexistence of three species through resource partitioning. To disentangle these possibilities, future studies should compare the coexistence patterns in homoxylan and xylose. For instance, Dal Bello et al. (2021) observed that the number of species coexisting in cellulose (a homopolysaccharide of glucose) was roughly double that of species coexisting in glucose (39). Thus, our results agree with the trend that complex substrates promote coexistence while simple substrates hinder it.

### Substrate complexity influences species interactions with more negative interactions in xylose than in xylan

Our study shows that in the polysaccharide xylan, pairs of species grew equally well in the presence of another species than in monoculture. In comparison, in the monosaccharide xylose, one of the species grew worse in the presence of another species than on its own. These pairwise dynamics scaled up to three species communities. For example, in xylose, species grew worse in the community than in monoculture. Our mathematical model showed that in xylose, these dynamics could be explained by resource competition, whereas the resource competition model could not predict the coexistence patterns in xylan.

One of the most interesting results is that, in xylan, all three species grew equally well in the community than alone. We posit that a combination of negative and positive interactions explains this observation (hypothesis H2c) rather than strict resource partitioning (hypothesis H2a). All three species encoded the genomic potential (endo-1-4-xylanases enzymes) to hydrolyze the xylose backbone to utilize the majority of the xylan polysaccharide (76% xylose). Given that we observed species compete for the monomer xylose (Figures 4B, 5B, 6B), we might expect species to compete and have reduced growth in the xylan community than in isolation, as predicted by the mathematical model (Figure 6A). However, we observed coexistence of the three species with a growth comparable in the communities than in isolation. Thus, we argue that a positive force, such as synergism, must counteract the negative interactions due to xylose competition. The combined effects of these positive and negative interactions would result in similar species’ densities in the community and in monoculture. Synergistic interactions on polysaccharides have been previously observed (40, 41). Even though in our system, we do not know the exact mechanism underlying positive interactions (if any), we speculate that it could be due to enzyme specialization. For example, *Flavobacterium* encodes unique copies of xylanases that the other species do not have (Table S2). Thus, the CAZymes in the community could synergize to hydrolyze xylan more completely (42).

In conclusion, we showed that interactions are less negative in the polysaccharide xylan than in the monosaccharide xylose. This is relevant because substrates in nature often exist in the form of polysaccharides, which can promote more positive interactions. Importantly, we have established a model community with three species from grass litter that stably coexist in the polysaccharide xylan. This synthetic community will be crucial for studies looking to understand the plant litter microbiome responses to climate change (43).

## Acknowledgments

We thank Melissa Martens for her assistance with the experiments. Special thanks to Jennifer Martiny for providing the strains and for providing valuable feedback on earlier manuscript versions. We thank all the collaborators from the Loma Ridge Climate Experiment, including Kathleen Treseder and Steve Allison, which are currently funded along with ARV by the National Science Foundation (award number 2308342). We thank Rodriguez-Verdugo lab members for providing useful feedback during this research. PA was supported by UC Irvine’s Summer Undergraduate Research Fellowship (SURP). ML was supported by the NIH-Maximizing Access to Research Careers (MARC) Grant. ARV was supported by UC Irvine’s School of Biological Sciences.

## Data Availability Statement

The genome sequences generated during the current study are available in the GenBank repository under accession numbers SAMN38146147 and SAMN38146146. All the other data generated or analyzed during this study are included in this published article and its supplementary information files.

## References

1. Coûteaux MM, Bottner P, Berg B. Litter decomposition, climate and liter quality. Trends Ecol Evol. 1995; 10:63–66.

2. Arnosti C, Wietz M, Brinkhoff T et al. The Biogeochemistry of Marine Polysaccharides: Sources, Inventories, and Bacterial Drivers of the Carbohydrate Cycle. Ann Rev Mar Sci. 2021; 13:81–108.

3. Bardgett RD, Freeman C, Ostle NJ. Microbial contributions to climate change through carbon cycle feedbacks. ISME J. 2008; 2:805–814.

4. Lindemann SR. A piece of the pie: engineering microbiomes by exploiting division of labor in complex polysaccharide consumption. Curr Opin Chem Eng. 2020; 30:96–102.

5. D’Souza GG, Povolo VR, Keegstra JM, Stocker R, Ackermann M. Nutrient complexity triggers transitions between solitary and colonial growth in bacterial populations. ISME J. 2021; 15:2614–2626.

6. Pontrelli S, Szabo R, Pollak S et al. Metabolic cross-feeding structures the assembly of polysaccharide degrading communities. Sci Adv. 2022; 8:eabk3076.

7. Tilman D. Resource competition and community structure. Monogr Popul Biol. 1982; 17:1–296.

8. Hardin G. The competitive exclusion principle. Science. 1960; 131:1292–1297.

9. Potts DL, Suding KN, Winston GC, Rocha AV, Goulden ML. Ecological effects of experimental drought and prescribed fire in a southern California coastal grassland. Journal of Arid Environments. 2012; 81:59–66.

10. Glassman SI, Weihe C, Li J et al. Decomposition responses to climate depend on microbial community composition. Proc Natl Acad Sci U S A. 2018; 115:11994–11999.

11. Martiny JB, Martiny AC, Weihe C et al. Microbial legacies alter decomposition in response to simulated global change. ISME J. 2017; 11:490–499.

12. Jansson JK, McClure R, Egbert RG. Soil microbiome engineering for sustainability in a changing environment. Nat Biotechnol. 2023; 41:1716–1728.

13. Berlemont R, Allison SD, Weihe C et al. Cellulolytic potential under environmental changes in microbial communities from grassland litter. Front Microbiol. 2014; 5:639.

14. Matulich KL, Weihe C, Allison SD et al. Temporal variation overshadows the response of leaf litter microbial communities to simulated global change. ISME J. 2015; 9:2477–2489.

15. Chase AB, Gomez-Lunar Z, Lopez AE et al. Emergence of soil bacterial ecotypes along a climate gradient. Environ Microbiol. 2018; 20:4112–4126.

16. Dolan KL, Peña J, Allison SD, Martiny JB. Phylogenetic conservation of substrate use specialization in leaf litter bacteria. PLoS One. 2017; 12:e0174472.

17. Chase AB, Arevalo P, Polz MF, Berlemont R, Martiny JB. Evidence for Ecological Flexibility in the Cosmopolitan Genus Curtobacterium. Front Microbiol. 2016; 7:1874.

18. Chase AB, Karaoz U, Brodie EL, Gomez-Lunar Z, Martiny AC, Martiny JBH. Microdiversity of an Abundant Terrestrial Bacterium Encompasses Extensive Variation in Ecologically Relevant Traits. mBio. 2017; 8:e01809–17.

19. Berlemont R, Martiny AC. Genomic potential for polysaccharide deconstruction in bacteria. Appl Environ Microbiol. 2015; 81:1513–1519.

20. Chase AB, Weihe C, Martiny JBH. Adaptive differentiation and rapid evolution of a soil bacterium along a climate gradient. Proc Natl Acad Sci U S A. 2021; 118:e2101254118

21. Bushnell B. A Fast, Accurate, Splice-Aware Aligner. 2014. https://www.osti.gov/biblio/1241166.

22. Bankevich A, Nurk S, Antipov D et al. SPAdes: a new genome assembly algorithm and its applications to single-cell sequencing. J Comput Biol. 2012; 19:455–477.

23. Chen Y, Ye W, Zhang Y, Xu Y. High speed BLASTN: an accelerated MegaBLAST search tool. Nucleic Acids Res. 2015; 43:7762–7768.

24. Langmead B, Salzberg SL. Fast gapped-read alignment with Bowtie 2. Nat Methods. 2012; 9:357–359.

25. Parks DH, Imelfort M, Skennerton CT, Hugenholtz P, Tyson GW. CheckM: assessing the quality of microbial genomes recovered from isolates, single cells, and metagenomes. Genome Res. 2015; 25:1043–1055.

26. Chaumeil PA, Mussig AJ, Hugenholtz P, Parks DH. GTDB-Tk: a toolkit to classify genomes with the Genome Taxonomy Database. Bioinformatics. 2019; 36:1925–1927.

27. Hyatt D, Chen GL, Locascio PF, Land ML, Larimer FW, Hauser LJ. Prodigal: prokaryotic gene recognition and translation initiation site identification. BMC Bioinformatics. 2010; 11:119.

28. Seemann T. Prokka: rapid prokaryotic genome annotation. Bioinformatics. 2014; 30:2068–2069.

29. Finn RD, Bateman A, Clements J et al. Pfam: the protein families database. Nucleic Acids Res. 2014; 42:D222–30.

30. Finn RD, Clements J, Eddy SR. HMMER web server: interactive sequence similarity searching. Nucleic Acids Res. 2011; 39:W29–37.

31. Berlemont R, Martiny AC. Phylogenetic distribution of potential cellulases in bacteria. Appl Environ Microbiol. 2013; 79:1545–1554.

32. Rodríguez-Verdugo A, Vulin C, Ackermann M. The rate of environmental fluctuations shapes ecological dynamics in a two-species microbial system. Ecol Lett 2019; 22(5):838–846.

33. Chang C-Y, Bajic D, Vila JCC, Estrela S, Sanchez A. Emergent coexistence in multispecies microbial communities. Science. 2023; 381:343–348.

34. Vandenberghe LPS, Karp SG, Pagnoncelli MGB et al. Chapter 2 - Classification of enzymes and catalytic properties. In: Singh SP, Pandey A, Singhania RR, Larroche C, Li Z, editors. Biomass, Biofuels, Biochemicals. Elsevier, 2020. p. 11–30.

35. Monod J. The growth of bacterial cultures. Annu Rev Microbiol. 1949; 3:371–394.

36. D’Souza GG, Schwartzman J, Keegstra J, Schreier JE. Interspecies interactions determine growth dynamics of biopolymer degrading populations in microbial communities. Proc Natl Acad Sci U S A. 2023; 120:e2305198120.

37. Daniels M, van Vliet S, Ackermann M. Changes in interactions over ecological time scales influence single-cell growth dynamics in a metabolically coupled marine microbial community. ISME J. 2023; 17:406–416.

38. Melo-Silveira RF, Viana RLS, Sabry DA et al. Antiproliferative xylan from corn cobs induces apoptosis in tumor cells. Carbohydr Polym. 2019; 210:245–253.

39. Dal Bello M, Lee H, Goyal A, Gore J. Resource-diversity relationships in bacterial communities reflect the network structure of microbial metabolism. Nat Ecol Evol. 2021; 5:1424–1434.

40. Deng YJ, Wang SY. Synergistic growth in bacteria depends on substrate complexity. J Microbiol. 2016; 54:23–30.

41. Deng YJ, Wang SY. Complex carbohydrates reduce the frequency of antagonistic interactions among bacteria degrading cellulose and xylan. FEMS Microbiol Lett. 2017; 364:fnx019

42. Malgas S, Mafa MS, Mkabayi L, Pletschke BI. A mini review of xylanolytic enzymes with regards to their synergistic interactions during hetero-xylan degradation. World J Microbiol Biotechnol. 2019; 35:187.

43. Rodríguez-Verdugo A. Evolving Interactions and Emergent Functions in Microbial Consortia. mSystems. 2021; 6:e0077421.

