## Supplementary Information for "Substrate complexity buffers negative interactions in a synthetic microbial community of leaf litter degraders"

Supplementary Figure 1

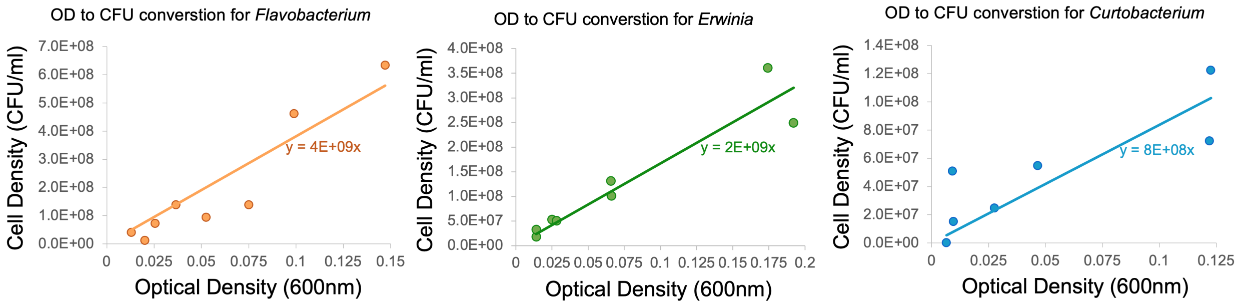


**Figure S1. Data used to convert the OD values to cell density values.** We used saturated cultures of *Flavobacterium* sp., *Erwinia* sp. and *Curtobacterium* sp., grown in M63 medium supplemented with xylose (16.67 mM) to relate the cell density (CFU/ml) to the optical density values. We fitted a linear relationship between the OD and the cell density (CFU/ml) and set the intercept to 0. The conversion rates for *Flavobacterium* sp., *Erwinia* sp. and *Curtobacterium* sp. were $4*10^9$, $2*10^9$ and $8*10^8$ CFU/ml, respectively.

Supplementary Figure 2

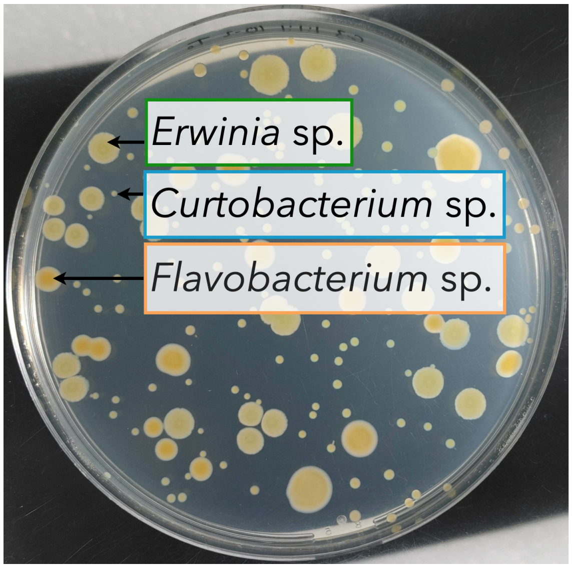


**Figure S2. The three species form different morphologies on a LB agar plate.** *Erwinia* sp. forms the largest colonies in LB agar, followed by *Flavobacterium* sp., which forms orange medium-size colonies. Finally, *Curtobacterium* sp. forms small colonies in LB agar.

Supplementary Figure 3

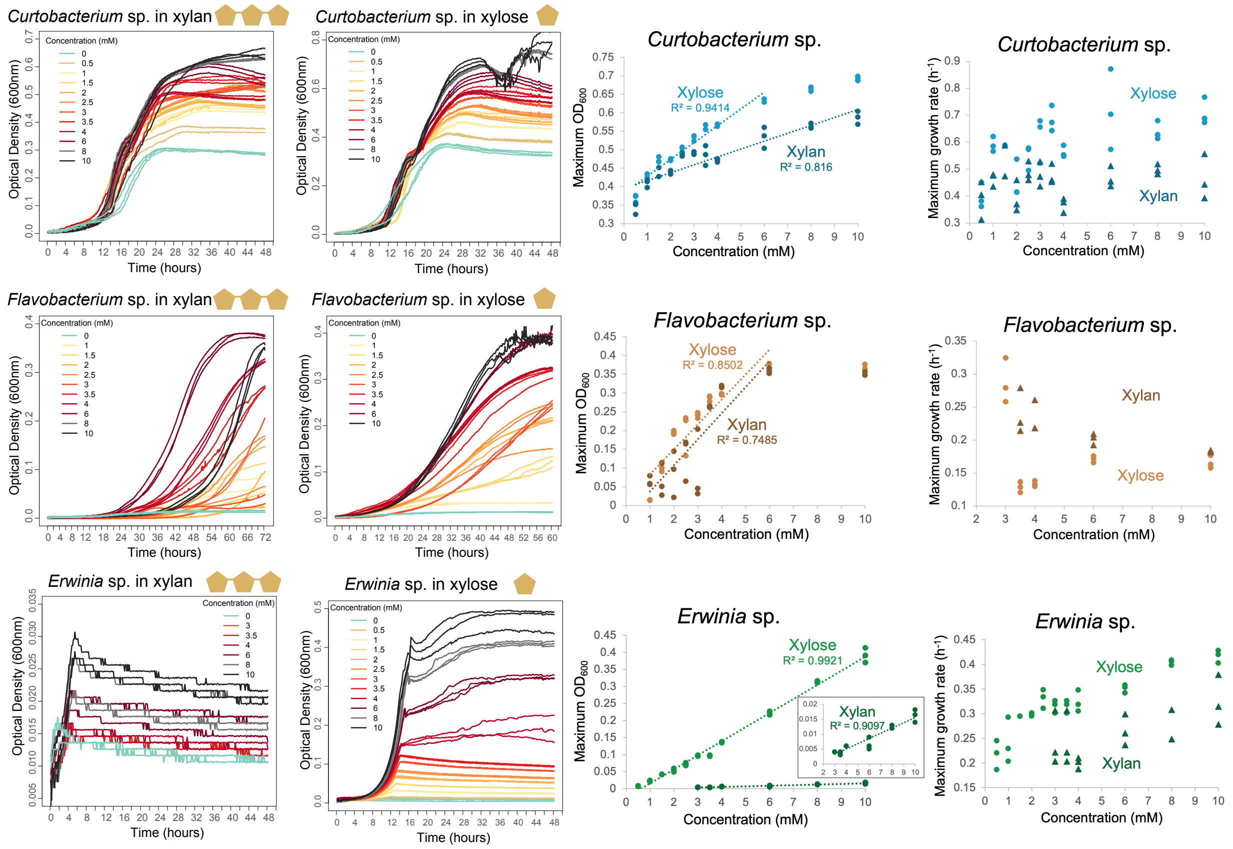


**Figure S3. Xylan and xylose are liming factors in the media**. Left panels: growth curves of species grown at different concentrations of xylan and xylose. The optical density measured at 600 nm over time is plotted for three technical replicates per concentration. Right panels: the linear relationship between the maximum population size and the concentration of xylose or xylan. Each dot corresponds to the maximum optical density ($OD_{600}$) achieved at the end of a growth cycle from the growth curves in the left panels. The dotted line is a linear regression fitting with its associated multiple R-squared.

#### Supplementary Figure 4

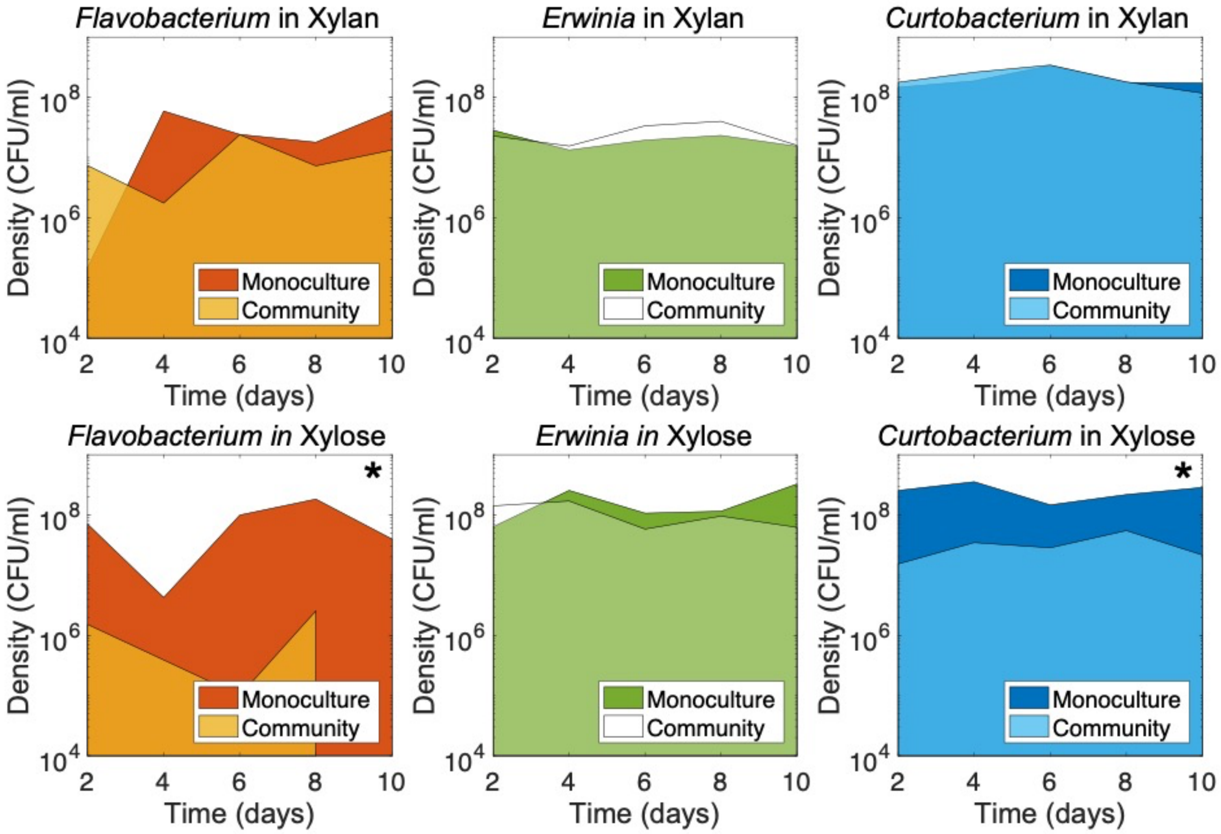


**Figure S4. Growth – estimated by the area under the curve – was not affected by other species in the community grown in xylan but was negatively impacted by other species in the community grown in xylose.** In xylan, the areas under the curve (AUC) of species grown in monoculture are similar to their AUCs in a community (top panels). Instead, in xylose, AUCs are lower in a three-species community than in a monoculture. *Flavobacterium* grew two orders of magnitude lower in the community than in monoculture in xylose ($4.983*10^6 \pm 2.786*10^6$ AUC in the community, $6.851*10^8 \pm 1.186*10^8$ AUC in monoculture; two-tailed t, $P = 0.029$). *Curtobacterium* grew one order of magnitude lower in the community than in monoculture in xylose ($2.733*10^8 \pm 1.032*10^8$ AUC in the community, $1.986*10^9 \pm 2.435*10^8$ AUC in monoculture; two-tailed t, $P = 0.010$). Each solid line corresponds to an average of three replicates of densities in monoculture and coculture. The area under the curve was estimated with the function `trapz` (Matlab version R2017a).

### Supplementary Figure 5

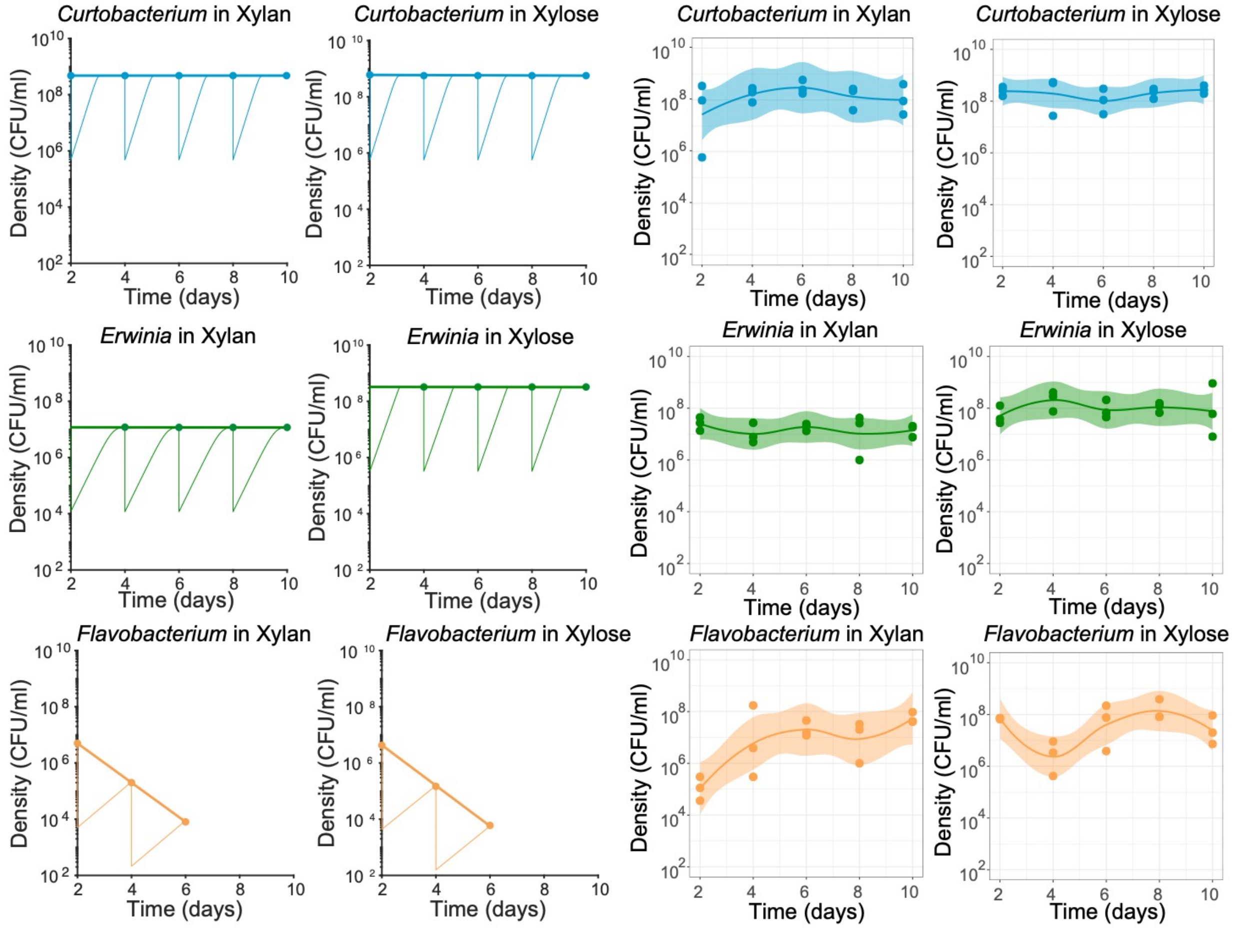


**Figure S5. The mathematical model recapitulates monocultures' dynamics for *Curtobacterium* sp. and *Erwinia* sp. but not for *Flavobacterium* sp.** On the left, the results derived from the mathematical model are plotted when simulating monocultures' growth during five serial transfers (1000-fold dilution every other day). Each tick line corresponds to the connection between the final densities of each species after 48h-cycles of growth, represented with thin lines. On the right are the experiment results (same as monocultures presented in Figure 2). Each line corresponds to a local polynomial regression fitting of three replicates with 95% confidence interval. Graphs were constructed with the function `stat_smooth` (R version 3.2.3).

**Table S1 Mathematical model**

| Bacterial growth | |
|---|---|
| *Curtobacterium*: | $\frac{dC}{dt} = \mu_{maxC} C \left(\frac{Xn}{k_{XnC} + Xn} + \frac{Xo}{k_{XoC} + Xo}\right)$ |
| *Flavobacterium*: | $\frac{dF}{dt} = \mu_{maxF} F \left(\frac{Xn}{k_{XnF} + Xn} + \frac{Xo}{k_{XoF} + Xo}\right)$ |
| *Erwinia*: | $\frac{dE}{dt} = \mu_{maxE} E \left(\frac{Xn}{k_{XnE} + Xn} + \frac{Xo}{k_{XoE} + Xo}\right)$ |
| **Resource changes** | |
| Xylan | $\frac{dXn}{dt} = -Xn\left(\frac{V_{XnC} Xn}{k_{XnC} + Xn} C + \frac{V_{XnF} Xn}{k_{XnF} + Xn} F + \frac{V_{XnE} Xn}{k_{XnE} + Xn} E\right)$ |
| Xylose | $\frac{dXo}{dt} = -Xo\left(\frac{V_{XnC} Xo}{k_{XoC} + Xo} C + \frac{V_{XoF} Xo}{k_{XoF} + Xo} F + \frac{V_{XoE} Xo}{k_{XoE} + Xo} E\right)$ |

| Variables | | Unit |
|---|---|---|
| $C$ | Density of *Curtobacterium* | unitless (OD) |
| $F$ | Density of *Flavobacterium* | unitless (OD) |
| $E$ | Density of *Erwinia* | unitless (OD) |
| $Xn$ | Concentration of xylan in culture | mM |
| $Xo$ | Concentration of xylose in culture | mM |
| **Parameters** | | **Unit** |
| $\mu_{maxC}$ | Maximum growth rate of *Curtobacterium* | $h^{-1}$ |
| $\mu_{maxF}$ | Maximum growth rate of *Flavobacterium* | $h^{-1}$ |
| $\mu_{maxE}$ | Maximum growth rate of *Erwinia* | $h^{-1}$ |
| $k_{XnC}$ | Half-saturation constant describing saturation of xylan by *Curtobacterium* | mM |
| $k_{XnF}$ | Half-saturation constant describing saturation of xylan by *Flavobacterium* | mM |
| $k_{XnE}$ | Half-saturation constant describing saturation of xylan by *Erwinia* | mM |
| $k_{XoC}$ | Half-saturation constant describing saturation of xylose by *Curtobacterium* | mM |
| $k_{XoF}$ | Half-saturation constant describing saturation of xylose by *Flavobacterium* | mM |
| $k_{XoE}$ | Half-saturation constant describing saturation of xylose by *Erwinia* | mM |
| $V_{XnC}$ | Maximum uptake rate of xylan by *Curtobacterium* | mM/unitless(OD)/h |
| $V_{XnF}$ | Maximum uptake rate of xylan by *Flavobacterium* | mM/unitless(OD)/h |
| $V_{XnE}$ | Maximum uptake rate of xylan by *Erwinia* | mM/unitless(OD)/h |
| $V_{XoC}$ | Maximum uptake rate of xylose by *Curtobacterium* | mM/unitless(OD)/h |
| $V_{XoF}$ | Maximum uptake rate of xylose by *Flavobacterium* | mM/unitless(OD)/h |
| $V_{XoE}$ | Maximum uptake rate of xylose by *Erwinia* | mM/unitless(OD)/h |

**Table S2. Genomic potential for xylan degradation.** A) Abundance of xylan-associated carbohydrate-binding modules (CBMs) and glycoside hydrolase (GH) families present in the genomes of the three strains used in this study. B) List of CBM /GH families associated with xylanase activity. Families are color-coded based on their catalytic properties (ref. 34). GH families in blue, cyan, and purple have a distinct catalytic domain with endo-1,4-ß-xylanase activity. GH families in yellow have residual or secondary xylanase activity.

**A)**

| **Strain** | **Total # CBM/GH** | **CBM9.1** | **CBM35** | **GH10** | **GH5** | **GH30** | **GH8** | **GH43** | **GH12** | **GH26** |
|---|---|---|---|---|---|---|---|---|---|---|
| *Curtobacterium* MMLR14002 | 71 | 0 | 2 | 1 | 0 | 0 | 1 | 1 | 0 | 1 |
| *Flavobacterium* LR40 | 199 | 3 | 1 | 2 | 7 | 3 | 2 | 13 | 1 | 2 |
| *Erwinia* LR17 | 38 | 0 | 0 | 0 | 0 | 0 | 1 | 1 | 0 | 0 |

**B)**

| **CBM/GH families** | **PfamID** | **Substrate; xylanase activity; gene** |
|---|---|---|
| CBM9.1 | PF06452 | cXylan; endo-1,4-ß-xylanase (EC3.2.1.8); *xynA* |
| CBM35 | PF16990 | cXylan; |
| GH10 | PF00331 | Xylan; endo-1,4-ß-xylanase (EC3.2.1.8), endo-1,3-ß-xylanase (EC3.2.1.32) |
| GH5 | PF00150 | Cellulose/Xylan; endo-1,4-ß-xylanase (EC3.2.1.8) |
| GH30 | PF02055 | Xylan; endo-1,4-ß-xylanase (EC3.2.1.8) |
| GH8 | PF01270 | Cellulose/Xylan; endo-1,4-ß-xylanase (EC3.2.1.8) |
| GH43 | PF04616 | Other Plant Polysaccharides; endo-1,4-ß-xylanase (EC3.2.1.8) |
| GH12 | PF01670 | Cellulose; residual or secondary xylanase activity |
| GH26 | PF02156 | Other Plant Polysaccharides; residual or secondary xylanase activity |
